# Insect—symbiont gene expression in the midgut bacteriocytes of a blood-sucking parasite

**DOI:** 10.1101/572495

**Authors:** Husnik Filip, Hypsa Vaclav, Darby Alistair

**Affiliations:** Biology Centre AS CR, Institute of Parasitology, Branišovská 31, České Budějovice 37005, Czech Republic; University of South Bohemia, Faculty of Science, Branišovská 31, České Budějovice 37005, Czech Republic; University of Liverpool, Institute of Integrative Biology, Crown Street, Liverpool L69 7ZB, UK

**Keywords:** RNA-Seq, B-vitamins, parasites, symbiotic bacteria, interactions, zinc, immunity

## Abstract

**Background:** Animals interact with a diverse array of both beneficial and detrimental microorganisms. These interactions sometimes spark obligate symbioses where the host depends on beneficial bacteria for survival and reproduction. In insects, these obligate symbioses in many cases allow feeding on nutritionally unbalanced diets such as plant sap and vertebrate blood. It is, however, still not clear how are these obligate intracellular symbioses maintained at the cellular level for up to several hundred million years. Exact mechanisms driving host-symbiont interactions are only understood for a handful of model species and data on blood-feeding hosts with intracellular bacteria are particularly scarce.

**Results:** Here, we analyzed interactions at the symbiotic interfaces of an obligately blood-sucking parasite of sheep, the louse fly *Melophagus ovinus*. We assembled a reference transcriptome from one male and one female individual and used RNA-Seq with five biological replicates to compare expression in the midgut cells housing bacteria to the rest of the gut (foregut-hindgut). We focused on nutritional and immunity interactions between the insect host and its obligate symbiont *Arsenophonus melophagi*, and also generated lower-coverage data for three facultative bacterial symbionts (*Sodalis melophagi, Bartonella melophagi*, and *Wolbachia* sp.) and one facultative eukaryote *Trypanosoma melophagium*. We found strong evidence for the importance of zinc in the system likely caused by symbionts using zinc-dependent proteases when acquiring amino acids, and for likely different immunity mechanisms controlling the symbionts than in closely related tsetse flies.

**Conclusions:** Our results show that cellular and nutritional interactions between this blood-sucking insect and its symbionts are less intimate than what was previously found in some (but not all) plant-sap sucking insects such as aphids, psyllids, whiteflies, and mealybugs. This finding is likely interconnected to several features observed in symbionts in blood-sucking arthropods, particularly their midgut intracellular localization (as opposed to being localized in truly specialized bacteriocytes), intracytoplasmic presence (as opposed to having an outermost host-derived ‘symbiosomal’ membrane), less severe genome reduction, and relatively recent associations caused by frequent evolutionary losses and replacements.

**Data deposition:** Raw RNA-Seq data were made available through the European Nucleotide Archive (ENA) database under the study accession number PRJEB30632. All assemblies and additional large supplementary files are available on FigShare [https://doi.org/10.6084/m9.figshare.6146777.v1]. All commands used for data analyses are available on Github [https://github.com/filip-husnik/melophagus].

## Background

Nutritional supplementation from symbiotic bacteria allowed several insect groups to become specialized on nutritionally unbalanced diets such as vertebrate blood and plant sap. Nutrients so far shown to be provided by bacteria include essential amino-acids and some B-vitamins in plant-sap sucking hosts, B-vitamins in blood-sucking hosts, and symbiotic bacteria (or protists and fungi) are also involved in nitrogen recycling in nitrogen-restricted environments such as in wood-feeding insects (Moran et al. 2008, McCutcheon and Moran 2011, Douglas 2016). What is still unclear is if there are any general insect-wide mechanisms driving these host-symbiont interactions. For example, how often do independent solutions of exactly the same functional problem emerge in even closely related species? To what extent do these interactions depend on the host biology (e.g. plant-sap vs. blood sucking insects) rather than on the host phylogeny (e.g. blood-sucking sucking Hemiptera vs. Diptera) or the symbiont phylogeny (e.g. Gammaproteobacteria vs. Bacteroidetes)? Since the small symbiont genomes are usually subsets of well-known bacterial genomes such as *E. coli* and code only a handful of unknown genes, it is feasible to roughly predict their metabolism either by simple mapping of present genes on metabolic pathways (Hansen and Moran 2013, Husnik and McCutcheon 2016) or by systems biology approaches such as Flux Balance Analysis (Thomas et al. 2009, MacDonald et al. 2011, Belda et al. 2012, Gonzalez-Domenech et al. 2012). What is not easily feasible, though, is to fully encompass and model the interactions with the host, especially for non-model species for which host genome, transcriptome or proteome data are not available.

### Our understanding of nutritional insect-symbiont provisioning comes mostly from plant-sap sucking insects

Majority of data concerning the host role in arthropod-bacteria symbiosis is undoubtedly available for pea aphids (Nakabachi et al. 2005, Gerardo and Wilson 2011, Hansen and Moran 2011, Poliakov et al. 2011). Hansen and Moran (2011) and Poliakov et al. (2011) untangled the intimate symbiotic interface in the pea aphid-*Buchnera* system, and confirmed the previously suggested (Nakabachi et al. 2005) host-symbiont cooperation in the production of essential amino acids (EAAs) and incorporation of ammonium nitrogen into glutamate by the GS/GOGAT (glutamine synthase/glutamine oxoglutarate aminotransferase) cycle. Flux-balance analyses and additional data (Macdonald et al. 2012) suggest that waste ammonia is recycled predominantly by the host cell (bacteriocyte) and that aphid aminotransferases (ornithine AT, branched-chain AT, and aspartate AT) incorporate ammonia-derived nitrogen into carbon skeletons synthesized by *Buchnera* to generate EAAs. These hypotheses were also verified experimentally (Macdonald et al. 2012). The highly similar picture observed in citrus mealybugs (Husnik et al. 2013), petiole gall psyllids (Sloan et al. 2014) or whiteflies (Luan et al. 2015), and identical enzymatic gaps in other endosymbiont genomes from hemipterans (Hansen and Moran 2013), imply that many (but perhaps not all; (Van Leuven et al. 2014)) insect hosts carry out these last steps to gain control of production of the final products. Unlike animals, plants can synthesize B-vitamins (Roje 2007), but whether B-vitamins are acquired by insects from the phloem/xylem sap of their host plants and provided to endosymbionts remains poorly understood. Endosymbiont genomes from plant-sap feeding insects retain several genes/pathways for biosynthesis of B-vitamins, e.g. biotin, riboflavin, and folate (Hansen and Moran 2013, Moran and Bennett 2014). It is unclear which B-vitamins are only used by symbionts and which are in addition also provided to their hosts. The only piece of experimental evidence implies that young symbiotic aphids are provided with riboflavin by their *Buchnera* endosymbionts (Nakabachi and Ishikawa 1999).

Host-symbiont cooperation is exceptionally dependent on the well-working transport of compounds between the bacteriocytes and the symbiont cells. Symbiotic bacteria of plant-sap sucking insects retain only a few general transporters, some of which very likely lost their substrate specificity (Charles et al. 2011). Host amino acid transporters can be involved in symbiont maintenance – some of them were extensively duplicated and specialized for bacteriocyte transfer (Duncan et al. 2014) and symbiont control (Price et al. 2013, Lu et al. 2016). No evidence of massive transfer of proteins among the symbiotic partners was so far confirmed, although one host protein was reported to be targeted to *Buchnera* cells in aphids (Nakabachi et al. 2014). However, such protein transfer is very likely needed in other hosts. For example, a recent rigorous analysis of host expression in two bacteriome types in a leafhopper host implies that nucleus-encoded genes usually supporting mitochondria also support bacterial endosymbionts (Mao et al. 2018).

### Nutritional interactions between blood-sucking insects and their symbiotic bacteria are understood only for a few hosts

Based on genomic data, different bacterial symbionts of blood-feeding insects can synthesize biotin, thiamine, riboflavin and FAD, panthotenate and coenzyme A, folate, pyridoxine, ubiquinol, nicotinamide, lipoic acid, and protoheme (Kirkness et al. 2010, Rio et al. 2012, Nikoh et al. 2014, Nováková et al. 2015, Boyd et al. 2016, Říhová et al. 2017). Controversy arises when discussing which particular cofactors are provided in particular host lineages. Interestingly, there are obligately blood-feeding arthropods (e.g. ticks or kissing bugs) that do not house stable intracellular microbes. These athropods either somehow efficiently extract rare nutrients from their blood diet or rely on extracellular gut bacteria acquired from the environment, e.g. by coprophagy as in kissing bugs (Eichler and Schaub 2002). Different blood-feeding lineages thus likely rely on symbionts for different subsets of these cofactors, perhaps due to differences in their blood-feeding strategies (e.g. facultative vs. obligate dependence on the blood diet), association with the host (temporary vs. permanent), host species (e.g. mammals vs. birds), enzymatic dependence (e.g. using alternative enzymes not depending on a particular co-factor), and evolutionary history. Some of the cofactors produced by symbionts are likely only used by symbiont-encoded enzymes rather than being provided to the insect host and other cofactors such as thiamine in human lice and louse flies are actually transported to the symbiont cells from the blood-diet with a thiamine ABC transporter encoded on the symbiont genome (reviewed in Husnik 2018).

In comparison to the very intimate host-symbiont cooperation on biosynthesis of amino acids in plant-feeding insects, bacteria in blood-feeding insects seem to be functioning more as independent units. The only RNA-Seq analysis from blood-feeding insects with intracellular symbionts was carried out in tsetse flies. The authors show that in terms of nutritional cooperation, only a few host genes seem to regulate and maintain the symbiosis, particularly a multi-vitamin transporter is up-regulated to shuttle B-vitamins from midgut bacteriocytes to haemolymph or other tissues (Bing et al. 2017).

### Insect immune response often distinguishes obligate mutualists from facultative symbionts and pathogens

Several ancient and intracellular obligate symbionts of insects have partially or completely lost bacterial cell envelope structures recognized by the insect immune system – peptidoglycan and lipopolysaccharides (McCutcheon and Moran 2011). In the latter case, they are often engulfed by a host-derived symbiosomal membrane (McCutcheon and Moran 2011), so there is nothing on their cell envelopes recognized as of bacterial origin by the host peptidoglycan-recognition proteins (PGRPs) or Gram-negative binding proteins (GNBPs). Interestingly, if there are some structures of bacterial origin still present as in lice symbionts, the hosts were either shown to jettison PGRPs, genes from the immunodeficiency signaling pathway (IMD), and many antimicrobial peptides (Gerardo et al. 2010, Kirkness et al. 2010) or modify them for symbiont defense. Blood-feeding insects that still keep PGRPs such as tsetse flies use amidase activity of one of PGRPs for peptidoglycan recycling in bacteriocytes (and milk glands of tsetse flies) and this activity shields symbionts from recognition by other PGRPs and expression of lineage-specific antimicrobial peptides mediated by the IMD (Anselme et al. 2006, 2008, Wang et al. 2009, Weiss et al. 2011, Wang and Aksoy 2012, Ratzka et al. 2013, Bing et al. 2017). A cautionary tale showing that insect symbioses are never-ending power struggles rather than idyllic interactions was reported from *Sitophilus* weevils and their recently ‘domesticated’ *Sodalis pierantonius* symbiont (Oakeson et al. 2014). A single antimicrobial peptide (Coleoptericin A) was shown to keep the symbionts under control, and if it was silenced by RNA interference, the symbionts were escaping from the bacteriocytes and spreading into host tissues (Login et al. 2011). The functional role of the symbionts is likely to synthesize a single non-essential amino acid, tyrosin, needed for cuticle hardness and when this benefit is no longer needed, symbiont numbers are reduced by the host (Vigneron et al. 2014). *Sodalis*-allied bacteria are very common symbionts of blood-sucking insects, so it is likely that similar symbiont control solutions had to also emerge in blood-feeding insects.

Here, we use comprehensive RNA-Seq data for an obligately blood-sucking insect harboring several endosymbionts and pathogens to understand metabolic interdependence and host-symbiont interactions. We generated data for an ectoparasite of sheep, *Melophagus ovinus*, and analyzed host-symbiont interactions within its microbiome containing not only the obligate symbiotic bacterium *Arsenophonus melophagi*, but also facultative bacteria *Sodalis melophagi, Bartonella melophagi*, and *Wolbachia* sp.; and the facultative eukaryote *Trypanosoma melophagium* (Small 2005, Chrudimský et al. 2012, Nováková et al. 2015, 2016).

## Results

### Contamination and rRNA depletion efficiency assessment

Since ribosomal RNA was depleted by a Terminator-5--Phosphate-Dependent Exonuclease treatment to allow parallel evaluation of expression for both the host and all its microbiome members, we tested the depletion efficiency by mapping trimmed reads against comprehensive 18S rRNA and 16S rRNA [small ribosomal subunit (SSU)] gene databases using PhyloFlash. Approximately 27-30 % of all reads per library mapped to these genes suggesting that the rRNA depletion method used had relatively poor efficiency. However, even if large subunit rRNA molecules (23S and 28S rRNA) make up more of the data than the small subunit rRNAs, total amount of sequencing data generated (371,432,771 trimmed reads) not only allowed us to assemble a high quality reference transcriptome for *M. ovinus*, but also to carry out abundance estimation and expression analyses for *M. ovinus* and *A. melophagi*. Species composition profiles allowed by SSU mapping revealed very homogenous microbiome composition across all samples (Figure 1A) with majority of data from the host *M. ovinus* (and low coverage sheep contamination from blood), its two most prominent bacterial symbionts *Arsenophonus* and *Sodalis melophagi*, and a eukaryotic commensal/pathogen *Trypanosoma melophagium.* Additional lower abundance taxa of interest include for example *Bartonella melophagi, Wolbachia* sp., and an unidentified ciliate related to sheep rumen ciliates. Species composition evaluation of the assembled metatranscriptome in Blobtools further corroborated these SSU results (Figure 1B).

**Fig 1.**
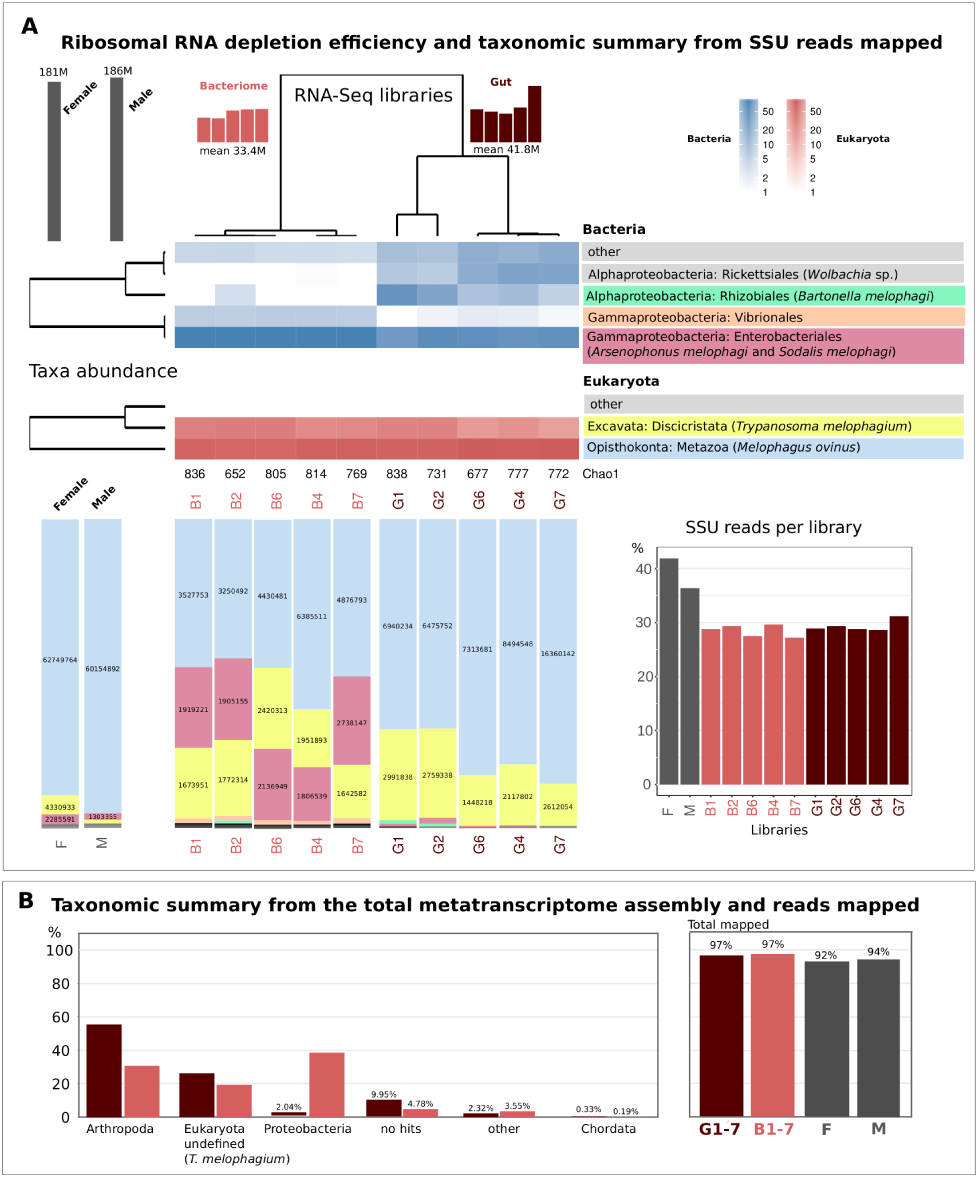
Ribosomal RNA depletion efficiency and species composition/contamination assessment. **(A)** Small ribosomal rRNA genes (both bacterial and eukaryotic) and **(B)** Total meta-transcriptome data. ‘Chao’ values represent total SSU diversity per library.

### Evaluation of symbiont expression

FPKM values obtained by mapping raw data on the *A. melophagi* genome are presented in Suppl. 1 and the improved *A. melophagi* annotation is available through FigShare (10.6084/m9.figshare.6146777). Figure 2 shows the expression values and predicted operon structure of the *A. melophagi* genome overlaid on its linear map with genes color-coded according to their COG (Clusters of Orthologous Groups) functional assignments. Expression values for the 20 most highly expressed *A. melophagi* genes are shown in Table 1.

**Fig 2.**
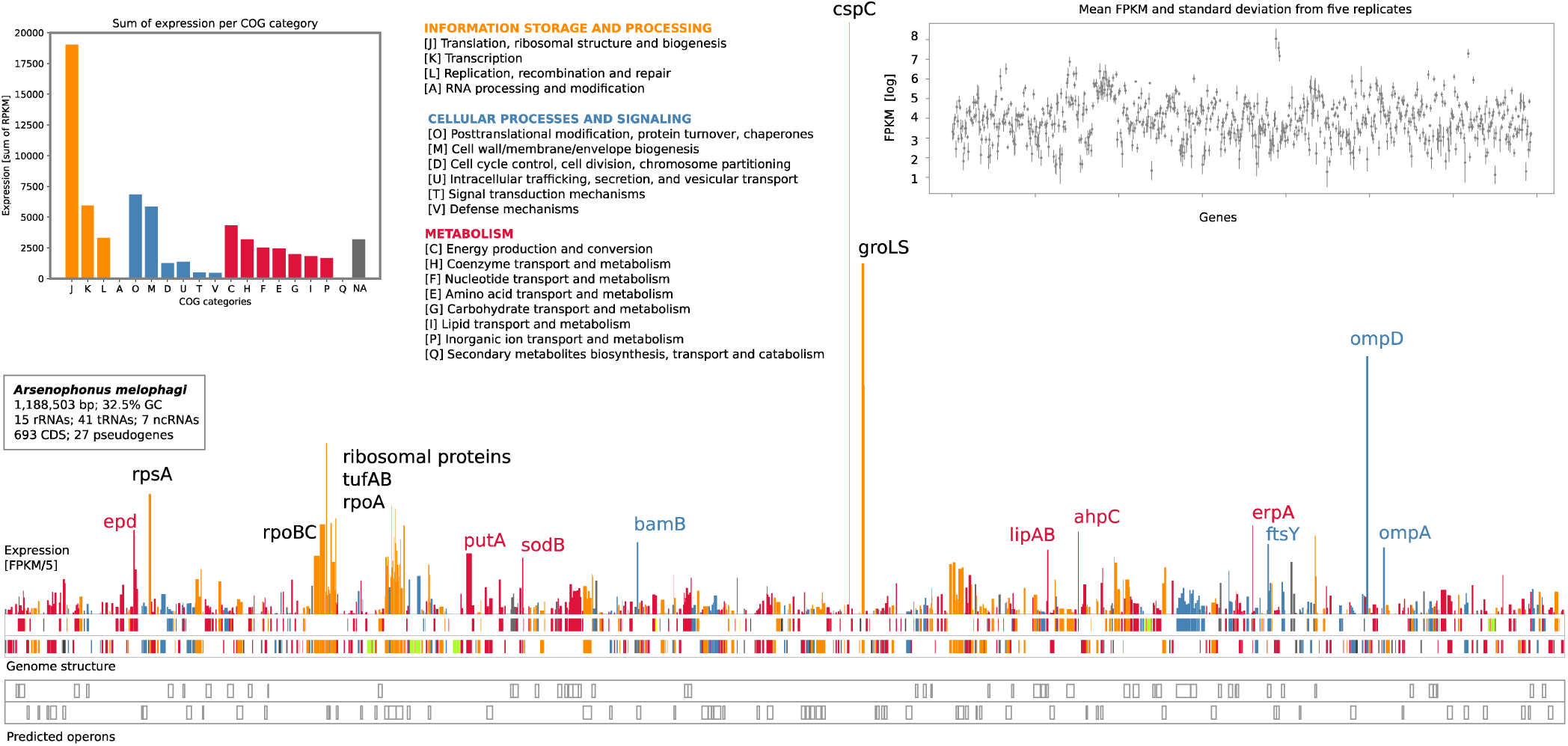
Linear genome map of *A. melophagi* genome overlaid by expression values of protein-coding genes. Genes are color-coded according to their broad COG functional categories: Information storage and processing in dark orange, Cellular processing and signaling in steel blue, and Metabolism in crimson. The most highly expressed genes (other than translation-related ribosomal proteins) are highlighted by gene names next to their respective peaks. Ribosomal RNA genes are shown in light green and tRNA/ncRNA genes in yellow. Expression of rRNAs is not shown since it was depleted and is likely biased. Expression of tRNAs and other short ncRNAs is not shown since these RNAs are under-represented due to RNA isolation and library preparation methods used. Top left inset: total sum of FPKM expression values for individual categories is included as a bar plot. Genes assigned to two or more categories were not included for simplicity. Top right inset: mean FPKM values with standard deviation showing consistent expression across five biological replicates of bacteriomes. The FPKM values are plotted on a log scale. Bottom: genes predicted to be expressed in operons.

**Tab 1:** Thirty most highly expressed *A. melophagi* genes. COG functional categories: [M] Cell wall/membrane/envelope biogenesis; [O] Post-translational modification, protein turnover, and chaperones; [U] Intracellular trafficking, secretion, and vesicular transport; [J] Translation, ribosomal structure and biogenesis [K] Transcription; [C] Energy production and conversion; [E] Amino acid transport and metabolism; [G] Carbohydrate transport and metabolism; [H] Coenzyme transport and metabolism; [P] Inorganic ion transport and metabolism. FPKM values are averaged from five replicates.

| Gene | Product | COG | FPKM |
| --- | --- | --- | --- |
| cspC | Cold shock-like protein | K | 3418 |
| groL | 60 kDa chaperonin | O | 2025 |
| ompD | Outer membrane porin | M | 1487 |
| groS | 10 kDa chaperonin | O | 1322 |
| rplJ | 50S ribosomal protein L10 | J | 986 |
| rpsA | 30S ribosomal protein S1 | J | 693 |
| rpmD | 50S ribosomal protein L30 | J | 628 |
| rplX | 50S ribosomal protein L24 | J | 609 |
| rpsL | 30S ribosomal protein S12 | J | 553 |
| tufA_1 | Elongation factor Tu | J | 524 |
| rpoB | RNA polymerase subunit $\beta$ | K | 518 |
| tufA_2 | Elongation factor Tu | J | 512 |
| erpA | Iron-sulfur cluster protein | C | 512 |
| epd | D-erythrose-4-P dehydrogenase | G | 486 |
| ahpC | Alkyl hydroperoxide reductase subunit C | O | 475 |
| rplM | 50S ribosomal protein L13 | J | 453 |
| rpsN | 30S ribosomal protein S14 | J | 447 |
| rplW | 50S ribosomal protein L23 | J | 430 |
| bamB | Outer membrane protein | M | 416 |
| rplN | 50S ribosomal protein L14 | J | 408 |
| ftsY | Signal recognition particle receptor | U | 405 |
| ompA | Outer membrane protein A | M | 386 |
| lipA | Lipoyl synthase | H | 371 |
| rplP | 50S ribosomal protein L16 | J | 365 |
| putA | Bifunctional protein PutA | E | 352 |
| rpoC | RNA polymerase subunit $\beta'$ | K | 336 |
| rpsM | 30S ribosomal protein S13 | J | 332 |
| rpsU | 30S ribosomal protein S21 | J | 330 |
| rpoA | RNA polymerase subunit $\alpha$ | K | 327 |
| sodB | Superoxide dismutase [Fe] | P | 325 |

### De-novo transcriptome assembly and differential expression analyses of host data

The transcriptomic data were de-novo assembled into a total of 166,038 transcripts (83,574 ‘genes’) with a contig N50 size of 1,456 bp. Total 166,522,714 bases were assembled. BUSCO completeness assessment resulted in 298 complete (202 duplicates due to the presence of *Trypanosoma melophagium* and sheep transcripts) and 5 fragmented genes out of 303 markers. Transdecoder ORF prediction resulted in 98,820 proteins (31,807 full-length; 35,448 partial; 31,565 internal) providing us with a robust proteome set for analysis of the host role in the symbiotic system. Our strictly decontaminated *M. ovinus* transcriptome (no transcripts with TPM < 1 or taxonomic assignment to Bacteria, Kinetoplastida or Chordata by Blobtools) consists of 43,315 transcripts with 23,780 predicted proteins (9,037 complete). These filtered data had the BUSCO score of 272 complete (80 duplicates), 8 fragmented, and 23 missing markers (out of 303 markers).

Raw counts of mapped reads, normalized digital expression values, and results of differential expression analyses from EdgeR for gut and 266 genes significantly down-regulated in the bacteriome tissues are available in Suppl. 2. In total, 1353 genes were found significantly up-regulated and 266 genes significantly down-regulated in the bacteriome section of the midgut (Suppl. 2). We note that the much larger number of up-regulated genes is an overestimate caused by bacterial transcripts that were not excluded from the total assembly for the differential expression step. The most highly up-/down-regulated genes that have functional annotation are shown in Table 2.

**Tab 2:**
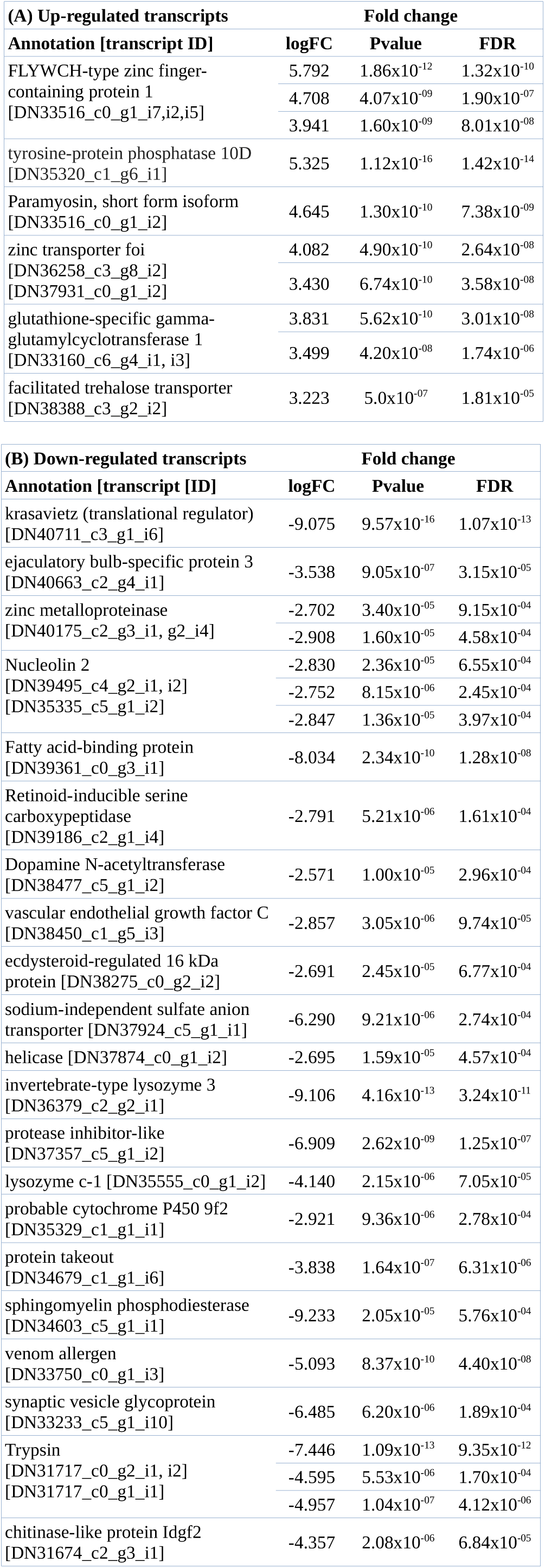
**(A)** Significantly up-regulated host transcripts in the bacteriome tissue. **(B)** Significantly down-regulated host transcripts in the bacteriome tissue (compared to the rest of the gut). We note that the most differentially expressed genes were uncharacterized proteins with no functional annotation – only genes with putative annotation are shown here in this table. At least four-fold expression change (logFC values shown) with p-value (Pvalue) cut-off for false discovery rate (FDR) set to 0.001 was required for a gene to be considered differentially expressed. Isoform TMM values equal or larger than 5.0 in at least one of the libraries were required for a gene to be included.

Interestingly, 11 transcripts significantly down-regulated in the bacteriome tissue come from unclassified RNA viruses (distantly related to the Hubei Diptera virus 14). Unfortunately, these transcripts are short and fragmented, perhaps due to their nucleotide diversity. Although an experimental verification of this finding is needed, it is tempting to speculate that either endosymbionts in the bacteriome tissue (*A. melophagi* and *S. melophagi*) protect from these viruses or that endosymbionts localized mostly outside the bacteriome tissue (*W. pipientis* and *B. melophagi*) promote the viral infection.

### Reconstruction of selected pathways involved in host —symbiont—pathogen interactions

B-vitamin and co-factor metabolism of all members of the microbiome set against the host pathways uncovered utilization of B-vitamins in the gut, acquisition of some B-vitamins (thiamine and pantothenate) by *A. melophagi* from bacteriocytes, but also likely indirect acquisition of other B-vitamins by the host bacteriocytes (e.g. after *A. melophagi* cell lysis). Possible exploitation of this resource by facultative members of the microbiome, *i.e. B. melophagi, S. melophagi, Wolbachia* sp., and *T. melophagium* is also likely (Fig 3B). However, the functional role of *T. melophagium* was not investigated in much detail as it likely does not regulate expression similarly to other trypanosomatids and it seems to be found along the entire gut (Fig 1). Gene expression data for *Sodalis melophagi and Bartonella melophagi* were of relatively low abundance, so we do not draw any strong conclusions from these data here. Host immune system reconstruction and identification of genes differentially expressed in the bacteriome and midgut sections provide evidence for a possible mechanism of how the host keeps its bacterial symbionts under control and how could facultative members of the microbiome escape its recognition (Fig 3A, Suppl 4). Gene/transcript information used for the reconstruction of the metabolic pathways and immune system can be found in Suppl. 1 and 2.

**Fig 3.**
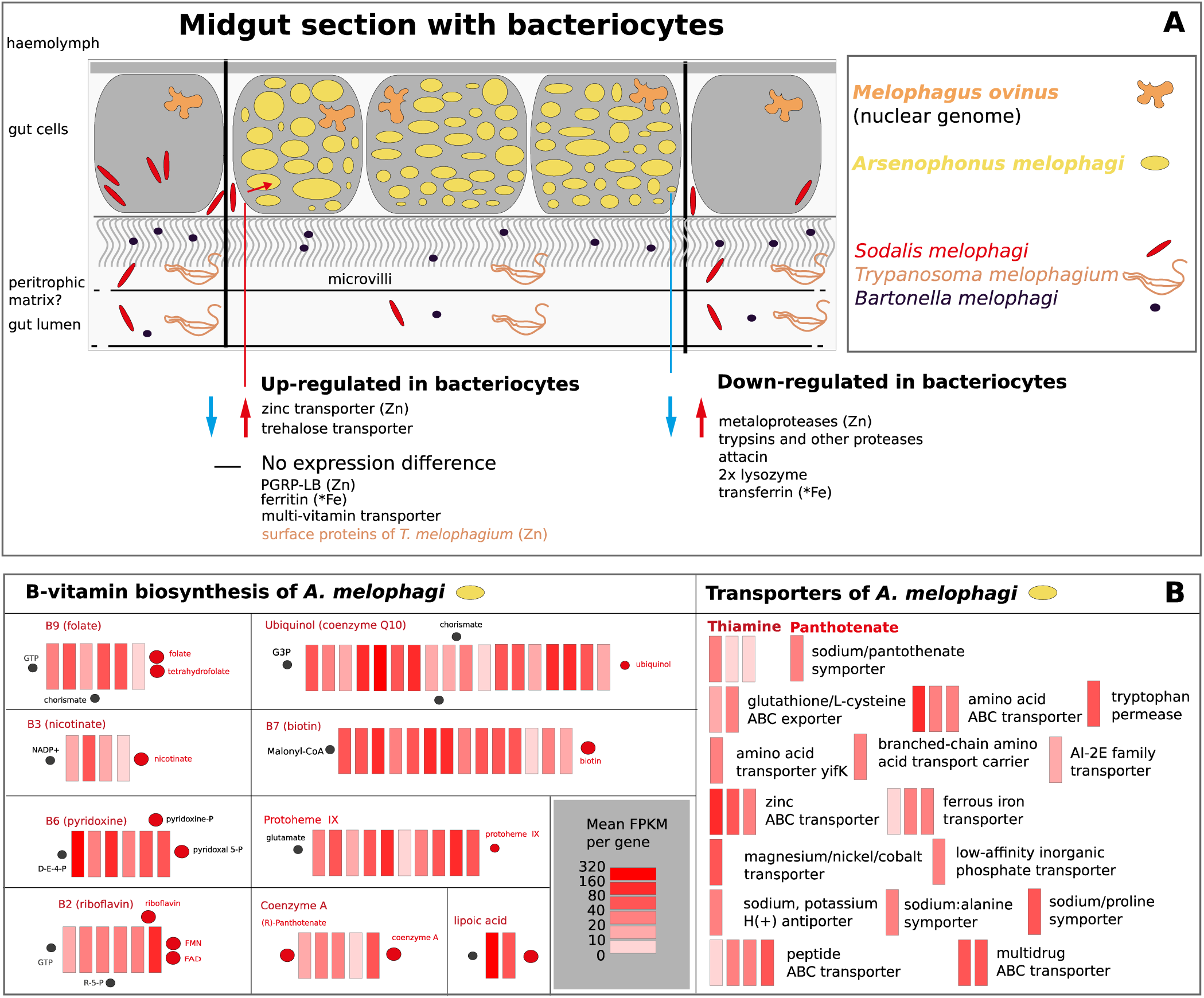
(A): Schematic reconstruction of nutritional and immunity processes revealed as involved in hos-symbiont interactions in the *M. ovinus* midgut cells harboring *Arsenophonus* symbionts. Selected host genes of interest up-regulated or down-regulated in bacteriocytes are highlighted. Zinc dependent enzymes are denoted by Zn, iron-binding proteins are denoted by Fe. **(B): B-vitamin biosynthetic pathways and transporters in *A. melophagi.*** Genes are filled by heatmap colors representing level of expression (mean FPKM from 5 replicates).

### Transporters up-regulated in bacteriocytes

Two host transporters possibly involved in maintenance of the symbiotic system were found to be significantly up-regulated in bacteriocytes (Tab 2). These transporters (facilitated trehalose transporter and zinc transporter) regulate transfer of zinc and trehalose to the cytoplasm of the bacteriocyte cells housing *A. melophagi.* Other host transporters found to be highly expressed along the entire gut include *e.g.* a sodium-dependent multivitamin transporter, a mitochondrial sodium-hydrogen exchanger, a sodium-potassium ATPase subunit alpha, a proton-coupled amino acid transporter, and a copper uptake protein. The expressed transporters of *A. melophagi* are highlighted in the Figure 3. Notably, thiamine and pantothenate transporters suggest that *A. melophagi* imports these two cofactors into its cells from the host blood diet or other microbiome members. Several transporters of *T. melophagium* were also evaluated as highly expressed in the bacteriome and/or gut sections (Suppl. 2). Trypanosomes thus likely scavenge available nutrients similarly as the remaining members of the microbiome: *S. melophagi, B. melophagi*, and *Wolbachia* sp. (Suppl. 2).

### Additional transcripts of bacterial origin

All transcripts recognized by our cut-off values as of bacterial origin are available in Suppl. 2. Most of these transcripts originated from endosymbionts present in the host species in both the gut and bacteriome tissues. The remaining bacterial transcripts represent low expression contamination by environmental and gut bacteria, false positive eukaryotic transcripts, or possible HGT candidates in the *M. ovinus* or *T. melophagium* genomes (Suppl. 2). No additional facultative symbiont species such as *Rickettsia* or *Cardinium* spp. were found to be present in our *M. ovinus* data by any of the methods employed (PhyloFlash, Blobtools, and BlastN/BlastX filtering). Since our reference genome assembly of *M. ovinus* (to double-check HGT candidates) is of extremely low quality, we do not discuss these genes of possible bacterial origin further. Genome data of much higher quality for both the host and its facultative microbiome members would be needed to reliably distinguish horizontal gene transfer events in the *Melophagus ovinus* genome and microbiome.

## Discussion

### Symbiont expression, B-vitamin biosynthesis, and host-symbiont metabolite exchange

The expression profile of *A. melophagi* (Fig 2; Suppl. 1; Tab 1) represents a typical example of an obligate symbiont in an intermediate stage of genome reduction. As expected for such a symbiont genome, genes for essential bacterial machinery (replication, transcription, translation) components such as ribosomal proteins, elongation factors, and RNA and DNA polymerases were among the most highly expressed. The most highly expressed gene was the cold shock-like protein CspC (and its anti-sense ncRNA) with transcription anti-terminator activity. Other highly expressed genes were found to be for heat shock proteins (GroEL and GroES), outer membrane proteins, and porins allowing various small solutes to cross the outer membrane (OmpD, OmpA, BamABDE), enzymes protecting against reactive oxygen species (alkyl hydroperoxide reductase and superoxide dismutase), and iron-sulfur cluster insertion protein ErpA; Fig 2; Suppl. 1). Very similar sets of highly expressed genes were also found in other obligate endosymbionts (Fukatsu and Ishikawa 1993, Aksoy 1995, Baumann et al. 1996, Fan et al. 2013). A pathway that shows unexpectedly high expression is conversion of proline to L-glutamate through the PutA enzyme and its subsequent conversion to D-glutamate by the MurI enzyme. Considering that proline is almost always the most common amino acid in insect haemolymph (Soulages 2010), it is likely that symbionts use proline-derived D-glutamate for their peptidoglycan cell wall. Proline is in insects generally stored as an energy reserve for energy-demanding activities (e.g. flight) or for adaptation to cold temperature, however, *M. ovinus* is flightless and permanently associated with its warm-blooded host. Its proline reserves can be thus potentially used by symbiotic bacteria for not only energy metabolism similarly to *Wigglesworthia* in tsetse flies (Michalkova et al. 2014), but proline can be also utilized for peptidoglycan synthesis. Moreover, proline is essential for trypanosome survival in tsetse flies (Mantilla et al. 2017) and could be also involved in triggering host–symbiont interactions as shown for example in *Xenorhabdus/Photorhabdus* symbionts (closely related to *Arsenophonus*) of nematodes (Crawford et al. 2010). Interestingly, B-vitamin and co-factor biosynthetic genes were not highly expressed in bacteriocytes with the exception of the lipoic acid pathway and several individual enzymes (Fig 2; Fig 3; Suppl. 1). This finding suggests that B-vitamins in blood-sucking insects, although essential, do not represent such a prominent case of host-symbiont cooperation as essential amino acids (EAAs) in plant-sap sucking insects. Such finding is not entirely unexpected since amino acids are crucial for proteosynthesis and growth of the organism, but vitamin-based co-factors are only needed by some of its enzymes. B-vitamins are based on our expression data likely used in relatively low volumes in adults and potentially needed most in particular host life stages that were not studied here (such as oocytes and developing larvae). At least two B-vitamins (thiamine and pantothenate) are acquired by *A. melophagi* from the blood and it is unknown if all the remaining B-vitamins that are actually synthesized are acquired by the host. An evolutionarily interesting case of B-vitamin biosynthesis needed for symbionts, but probably not provided to the host, was found in cicadas. Obligate co-symbiont of cicadas, *Hodgkinia cicadicola*, retained cobalamine-dependent methionine biosynthesis, so it has to devote at least 7% of its proteome to synthesize cobalamine – vitamin B12 (McCutcheon et al. 2009). Future experimental studies will need to be conducted to determine whether a similar scenario also applies to some pathways in blood-sucking insects.

We saw only a few host transporters up-regulated specifically in the bacteriocytes of *M. ovinus*, as would be expected in a system without a symbiosomal membrane and with an obligate symbiont still retaining a decent set of both specific and non-specific transporters (Fig 3; Suppl. 2). Because transporters likely import essential nutrients from the gut lumen to the endothelial cells along the whole gut, we are unable to fully recognize which of these highly expressed transporters transport the nutrients utilized also by symbionts. Two transporters clearly up-regulated in bacteriocytes (facilitated trehalose transporter and zinc transporters; Suppl. 2) support the symbiotic association by providing the metabolites to be processed either by the host bacteriocyte cell or the symbiont cells. We note that the sodium-dependent multi-vitamin transporter is not significantly up-regulated in bacteriocytes (contrary to the situation in tsetse flies), but rather highly expressed along the entire gut.

Several hypotheses can be put forward concerning the cell specialization of midgut bacteriocytes. First, these gut cells could be quite polarized and therefore able to carry out a number of different functions depending on their localization. For example, different transporters are likely to be expressed on the gut lumen side compared to the haemolymph side. Our data are averaged from the total tissue and do not allow distinguishing such subtle expression differences that would be only detectable with single-cell RNA-Seq. An alternative explanation is that these cells are so tightly packed with endosymbionts that they are no longer able to function as regular midgut cells and are thus acting as a specialized space for harboring endosymbionts, with only a few essential host genes up-regulated. Since these two scenarios are not mutually exclusive, we recognize that the outcome is likely a gene/pathway-specific trade off between these two (and likely a few additional) explanations.

### Metal metabolism – comparison to tsetse flies and implications for blood digestion

Iron regulation (*i.e.* acquisition, utilization, storage, and transport) is an essential function for blood-sucking insects needed to avoid iron toxicity. It is usually carried out by two iron binding proteins (IBP) – ferritin and transferrin. Similar to tsetse flies, heavy and light chain transcripts of ferritin are highly expressed ubiquitously along the whole gut, confirming their general role in iron transport and storage (Strickler-Dinglasan et al. 2006). Particularly, ferritin is involved in sequestering iron from a blood meal to avoid oxidative stress. Transferrin was down-regulated 4.5-fold in bacteriocytes, suggesting that its role is much more specific and perhaps connected to regulation of microorganisms in the gut. It was shown that it can also act as an antimicrobial protein sequestering iron from pathogens (Yoshiga et al. 1999). *A. melophagi* does not retain its own ferritin like *W. glossinidia*, but codes and expresses the iron transporter genes feoABC and the ferric transcriptional regulator (fur).

Surprisingly, multiple lines of evidence support a significance of zinc for this obligate symbiosis. Zinc is generally the second most abundant metal (after iron) in most organisms. It is essential for hundreds of enzymes and zinc finger-containing transcription factors (Coleman 1992, 1998, McCall et al. 2000). Several transcripts for zinc transporters are highly up-regulated in bacteriocytes and very lowly expressed in the gut (Tab 2). Furthermore, the zinc ABC transporter of *A. melophagi* symbionts is highly expressed (Fig 3AB; Suppl. 1) and thus supports that zinc is not only imported into the cytoplasm of bacteriocytes, but also imported into *A. melophagi* cells. Host zinc proteases are likely enzymes predominantly using zinc in the gut and two of them were shown to be highly expressed in the gut of tsetse flies (Yan et al. 2002). In our study, many zinc proteases were highly expressed in the rest of the gut, but down-regulated in bacteriocytes (Tab 2; Suppl. 2), suggesting that zinc is mostly used by symbionts in the bacteriome section of the midgut.

A question of particular importance is which *A. melophagi* enzymes are responsible for this zinc dependence on their host and how symbionts participate on the blood digestion. There are numerous zinc-dependent enzyme candidates obvious from our data (Suppl. 1), particularly several zinc proteases (TldD, TldE, HtpX, FtsH, RseP, YebA) and a putative metalo-beta-lactamase (GloB). Zinc proteases are involved in proteolysis, suggesting that *A. melophagi* symbionts acquire amino acids by digesting peptide bonds of proteins. This finding could explain why aposymbiotic tsetse flies have difficulties digesting blood (Pais et al. 2008). Beta-lactamases are enzymes that provide bacteria with resistance to beta-lactam antibiotics such as penicillin, ampicillin, and many others. Metalo-beta-lactamases in particular are well-known for their resistance to a broad spectrum of beta-lactam antibiotics and beta-lactamase inhibitors (Bradford 2001, Drawz and Bonomo 2010). Since sheep in the Czech Republic are frequently treated with beta-lactam antibiotics to avoid bacterial infections, it is tempting to speculate that *Arsenophonus* symbionts use their metalo-beta-lactamase (and *Melophagus* supports them by providing zinc) to avoid elimination by antibiotics circulating in sheep blood. If confirmed by additional experiments, it would be, to our knowledge, the first case of an obligate endosymbiont retaining antimicrobial resistance (AMR) genes to survive in its blood-sucking host frequently exposed to antibiotics.

### Insect immunity, peritrophic matrix, unknown proteins, and the role of gene specialization

Among the transcripts up-regulated in the whole gut and down-regulated in bacteriocytes (Suppl. 2), the antimicrobial peptide attacin (−3.3 FC) and two lysozymes (−550.9 FC and −17.6 FC) are very likely responsible for controlling bacterial infections in the gut and not targeting obligate symbionts *A. melophagi* (Fig 3A). However, their exact role (if any) in the control of *S. melophagi* and *B. melophagi* remains to be elucidated, although both can be highly effective on environmental bacteria. Strikingly, PGRP-LB, although highly expressed in both tissues, is not up-regulated in bacteriocytes (Fig 3A, Suppl. 3) as was reported for the tsetse fly system (Wang et al. 2009, Bing et al. 2017). This finding implies that the host likely unselectively recycles peptidoglycan along the whole gut and, since GNBP has very low expression along the gut, it is unknown if there is an additional mechanism targeting environmental infections in specific sections of the midgut. Possible implications of this difference might be either that the host does not target *S. melophagi* and *B. melophagi*, but is currently ‘domesticating’ these bacteria because they can increase its fitness (e.g. by providing thiamine or pantothenate not synthesized but needed by *Arsenophonus*), or that the host has evolved as partially immunocompromised because of its relatively bacteria-free diet. The latter explanation would be supported by the lack of peritrofic matrix in *M. ovinus* reported by early microscopy studies (Waterhouse 1953, Lehane 1997). One of essential functions of this non-cellular membrane that is generally found in all other Hippoboscidae species (Waterhouse 1953, Lehane 1997) is providing a barrier to infection by pathogens. Our RNA-Seq data do not include chitin synthesis genes of insect origin expressed in the gut, corroborating the microscopy observations.

As usual in insect RNA-Seq studies, many differentially expressed and/or highly expressed genes observed in our data (Suppl. 2) code conserved proteins with unknown function, hypothetical proteins, or orphan proteins with no or too low homology to proteins in protein databases. At least some of these genes are very likely involved in symbiosis maintenance and need to be examined experimentally in future studies. For example, RNA-Seq study on aphid bacteriocytes and whole mount *in situ* hybridizations of over-represented transcripts encoding aphid-specific orphan proteins has revealed a novel family of small cysteine-rich proteins with signal peptides (Shigenobu and Stern 2013). This finding supports hypotheses that numerous orphan genes in the pea aphid genome (International Aphid Genomics Consortium 2010) evolved to assist in lineage specific traits, such as symbiosis. A very similar scenario can be possibly applied to other symbiotic insects. The role of gene or even isoform specialization and recruitment for functioning in bacteriocytes was observed also in our data, for example in zinc transporters and proteinases (Suppl. 2).

### Host-symbiont interactions in blood-sucking and plant-sap sucking insects – similarities and differences

Unlike in plant-sap sucking insects, where the host is strongly involved in the biosynthesis of EAAs by providing the starting products and carrying out the last aminotransferase steps of the pathways, we found no evidence for such intimate interactions in our *M. ovinus* data for the B-vitamin or other biosynthetic pathways (Tab 1; Fig 3; Suppl. 2). Whether this is a common situation in all blood-sucking insects, or if it applies only to hosts with symbionts having medium-sized genomes, remains to be investigated. As insect symbionts are either present directly in the bacteriocyte cytoplasm (common in blood-sucking and omnivorous hosts), or enclosed in a so-called symbiosomal membrane (common in plant-sap sucking insects), the cellular localization of symbionts may also be responsible for the intimacy of the relationships. The cytoplasm-harbored symbionts, such as *A. melophagi* in *M. ovinus*, have direct access to all nutrients available in the host cells. For symbionts covered by a symbiosomal membrane, their host controls which nutrients will be available to the symbionts by adjusting the expression of transporters through the symbiosomal membrane localization (Price et al. 2013, Duncan et al. 2014). This situation then inevitably leads to highly interconnected relationships. Moreover, if the bacteriome section is localized in the gut (e.g. in tsetse flies, louse flies, and carpenter ants), it cannot be fully specialized for interactions with the symbionts, because it still has to express at least a handful of genes for digestion of its food source. Evaluating the exact impact of this less intimate symbiotic integration can eventually help to untangle the gradual steps in genome reduction observed for endosymbiont genomes.

## Conclusions

Using RNA-Seq with five biological replicates of the midgut cells housing bacteria and the rest of the gut, we uncovered the interactions among a blood-sucking parasite and its microbiome. We found strong evidence for importance of zinc in the system possibly caused by participation of symbionts on blood-digestion; and for different immunity mechanisms controlling symbionts than in closely related tsetse flies. Our results show that this blood-sucking insect is much less intimately involved in cooperation on biosynthesis of nutrients than plant-sap sucking insects such as aphids, psyllids, whiteflies, and mealybugs. This finding is likely interconnected to several features observed in symbionts in blood-sucking arthropods, particularly intracytoplasmic midgut localization of bacteria, their less severe genome reduction, and younger associations caused by frequent evolutionary losses and replacements. Since the specialized bacteriome section of *M. ovinus* is clearly distinct from the rest of the midgut based on both morphology and gene expression, it will be interesting to compare gene expression of this specialized tissue to other blood-sucking insects in future.

## Methods

### RNA extraction and sequencing

*Melophagus ovinus* parasites were sampled from the same population as used for our previous studies (Chrudimský et al. 2012, Nováková et al. 2015). Insects were immediately dissected in RNAlater (Qiagen) to stabilize expression profiles and kept deeply frozen until RNA extractions. Five biological replicates of bacteriomes and five replicates of the remaining portion of gut from identical samples were prepared for RNA-Seq. Total RNA was extracted from pools of five individual bacteriomes and guts for seven replicates using RNeasy Mini Kit (Qiagen). Five of the replicates for both tissues were selected for sequencing based on RNA quality and Bioanalyzer chip results (Agilent). All samples were DNased by RNase-free DNase I (Ambion) and ribosomal RNA was depleted by a Terminator-5--Phosphate-Dependent Exonuclease treatment (Epicentre). Ten RNA-Seq libraries were prepared from the enriched RNA samples using the ScriptSeq strand-specific protocol (Epicentre). Paired-end sequencing (2×100 bp) of the ten RNA-Seq libraries was multiplexed on one lane of the Illumina HiSeq 2000 platform at the Centre for Genomic Research, University of Liverpool. Total RNA was also extracted from one whole female and one whole male *M. ovinus* to improve *de-novo* assembly of the reference transcriptome and generate a sufficient gene set. The identical procedure as described above was carried out and the two libraries were multiplexed on one lane of the HiSeq 2000. Raw Fastq files were trimmed for the presence of Illumina adapter sequences using Cutadapt v1.1 [http://code.google.com/p/cutadapt]. Option -O 3 was used, so that the 3’ end of any read which matched the adapter sequence for 3 bp or more was trimmed. The reads were further quality-trimmed by Sickle v1.200 [https://github.com/najoshi/sickle] with a minimum window quality score of 20. Reads shorter than 10 bp after trimming and singlet reads were removed. The quality-trimming resulted in 371,432,771 read pairs. PhyloFlash v3.1 (Gruber-Vodicka et al. 2019) was used for rRNA depletion efficiency and contamination assessment using 18S rRNA and 16S rRNA gene databases.

### Arsenophonus melophagi gene expression and analyses

Bacterial expression was analyzed by mapping reads from individual libraries to the *Arsenophonus melophagi* assembly with Bowtie2 (Langmead and Salzberg 2012). Bam files were imported into BamView integrated in the Artemis browser v15.1.1 (Carver et al. 2012). The *A. melophagi* annotation was subsequently improved according to the expression data, i.e. mainly pseudogene remnants (not-expressed short hypothetical proteins) were re-annotated. These pseudogene re-annotation results were also supported by our custom pipeline [https://github.com/filip-husnik/pseudo-finder]. Identification of transcript boundaries, quantification of transcript abundance (FPKM values), and prediction of operon structure was carried out in Rockhopper v2.0.3 (McClure et al.2013).

### De-novo metatranscriptome assembly and differential expression analyses

De-novo metatranscriptome assembly was carried out by the Trinity assembler v2.4.0 (Grabherr et al. 2011) from digitally normalized read pairs (targeted maximum coverage set to 50) using strand-specific information. RSEM 1.3.0 (Li and Dewey 2011) was used to count mapped reads. The trimmed mean of M-values normalization (TMM), generation of normalized TPM values (transcripts per million transcripts), and differential expression analyses with five biological replicates were carried out in EdgeR (Robinson et al. 2010) BioConductor package. Only transcripts with at least four-fold expression change (p-value cut-off for false discovery rate set to 0.001) were considered to be differentially expressed between the bacteriome and gut tissues.

### Reference transcriptome filtering for bacterial and eukaryotic contamination

The reference transcriptome of *M. ovinus* was filtered for likely assembly artifacts and lowly supported transcripts so that only transcripts with normalized TPM >1 in at least one sample or replicate were retained (retained 30.95%, i.e. 51,386 from 166,038 of total transcripts). Taxonomy was assigned to all transcripts by Blobtools (Laetsch and Blaxter 2017) with the ‘--bestsum’ flag based on Blastn searches against the NT database and Diamond BlastX against the Uniprot proteome database. All transcripts assigned either to the superkingdom Bacteria (symbionts and bacterial contamination), the phylum Chordata (sheep and human contamination) or the Kinetoplastida class (*Trypanosoma melophagium*) were removed from the reference transcriptome. However, we note that some of *Trypanosoma melophagium* transcripts could be missed by this approach since transcripts with no hits were retained. Single-gene phylogenetic inference would be needed to fully decontaminate *T. melophagium* contamination from the *Melophagus* reference transcriptome since no reference genome exists for neither *M. ovinus* nor *T. melophagium*. Since no high-quality genome data are available for *M. ovinus, T. melophagium, S. melophagi*, and *B. melophagi*, we did not inspect further the bin of all transcripts filtered out as bacteria (e.g. to detect possible horizontal gene transfers).

### Transcriptome annotation and protein prediction

TransDecoder [https://github.com/TransDecoder/] was used for ORF and protein prediction of the filtered reference proteome. Complete functional annotation was produced by Trinotate and final results were uploaded into a custom MySQL database and analyzed through TrinotateWeb (Grabherr et al. 2011). BUSCO v3 was used to asses the transcriptome completeness against 303 universally conserved eukaryotic proteins (Simão et al. 2015) both before and after contamination removal.

### Reconstruction of selected pathways, metabolite exchange, and immune system composition

Digital expression values were overlaid on *A. melophagi* pathway map in the Pathway Tools Software (Karp et al. 2010). Up-regulated host transcripts of interest (e.g. B-vitamin, co-factor, metal, and amino acid metabolism) were analyzed manually for possible interactions with symbionts. Previously published B-vitamin pathways were updated with this expression information. Expression of genes annotated in the *A. melophagi* genome as transporters was assessed to reconstruct metabolite flux to and from the symbionts. Similarly, annotations of up-regulated host transcripts were screened for transporters to analyze the host role in the maintenance of symbiotic tissue. Transporter candidates were checked by BlastP (e-value 1e^-6^) against the NR protein database. EMBL-EBI CoFactor database (Fischer et al. 2010) was used for searches of co-factor enzyme dependence.

The insect immune system was reconstructed by blast searches (BlastP, BlastN, and BlastX) of homologs against a custom blast database built from our transcriptome assembly and gene candidates were tested against the non-redundant NCBI database. In particular, *Drosophila melanogaster* and *Glossina morsitans* homologs acquired from the Insect Innate Immunity Database (Brucker et al. 2012) and VectorBase (https://www.vectorbase.org/) were used as queries. Pathways involved in control of symbiotic bacteria in gut and bacteriome tissues were compared to the situation in tsetse flies using literature review.

## Description of additional data files

**Suppl. 1** Table with symbiont FPKM values for individual replicates, mean FPKM values, COG categories, locus tags, gene names, and protein products for the *A. melophagi* genome.

**Suppl. 2** Table with raw counts of mapped reads, normalized FPKM values, and EdgeR results for *M. ovinus*.

**Suppl. 3** Figure and table with IMD, JAK/STAT, JNK, and Toll genes detected in the *M. ovinus* transcriptome.

Additional large supplementary files are available on FigShare [https://doi.org/10.6084/m9.figshare.6146777.v1]. In particular, the improved *A. melophagi* genome annotation and pathway maps with highlighted expression, *M. ovinus* transcriptome annotation report, total meta-transcriptome assembly (including predicted proteins), filtered reference transcriptome (including predicted proteins), complete results of the Trinity-Transdecoder-RSEM-edgeR-Trinotate pipeline (as a TrinotateWeb MySQL database), and supplementary figures with quality check analyses for individual samples/replicates and differential expression visualization (Volcano and MA plots).

## Author’s contributions

FH designed and coordinated the study, carried out the preparation of samples, analyses of data, and drafted the manuscript. VH carried out analyses of immune genes and participated in the study design and manuscript revisions. AD designed the study and participated in manuscript revisions. All authors read and approved the final manuscript.

## Acknowledgements

We thank the Centre for Genomic Research (University of Liverpool) for sequencing services. FH was supported by a postdoctoral research fellowship from EMBO (ALTF 1260-2016) while writing this article.

## List of abbreviations used

COG: Clusters of Orthologous Groups
EAAs: Essential Amino Acids
FPKM: Fragments Per Kilobase Million
GNBP: Gram-Negative Binding Protein
HGT: Horizontal Gene Transfer
TPM: Transcripts Per Kilobase Million
TMM: Trimmed Mean of M-values normalization method
NR and NT: Non-redundant protein and nucleotide databases
ORF: Open Reading Frame
RNA-Seq: RNA Sequencing
PGRP: Peptidoglycan Recognition Protein
IMD: Immunodeficiency signaling pathway

