## Supplementary material for "Insect—symbiont gene expression in the midgut bacteriocytes of a blood-sucking parasite": Suppl. 3

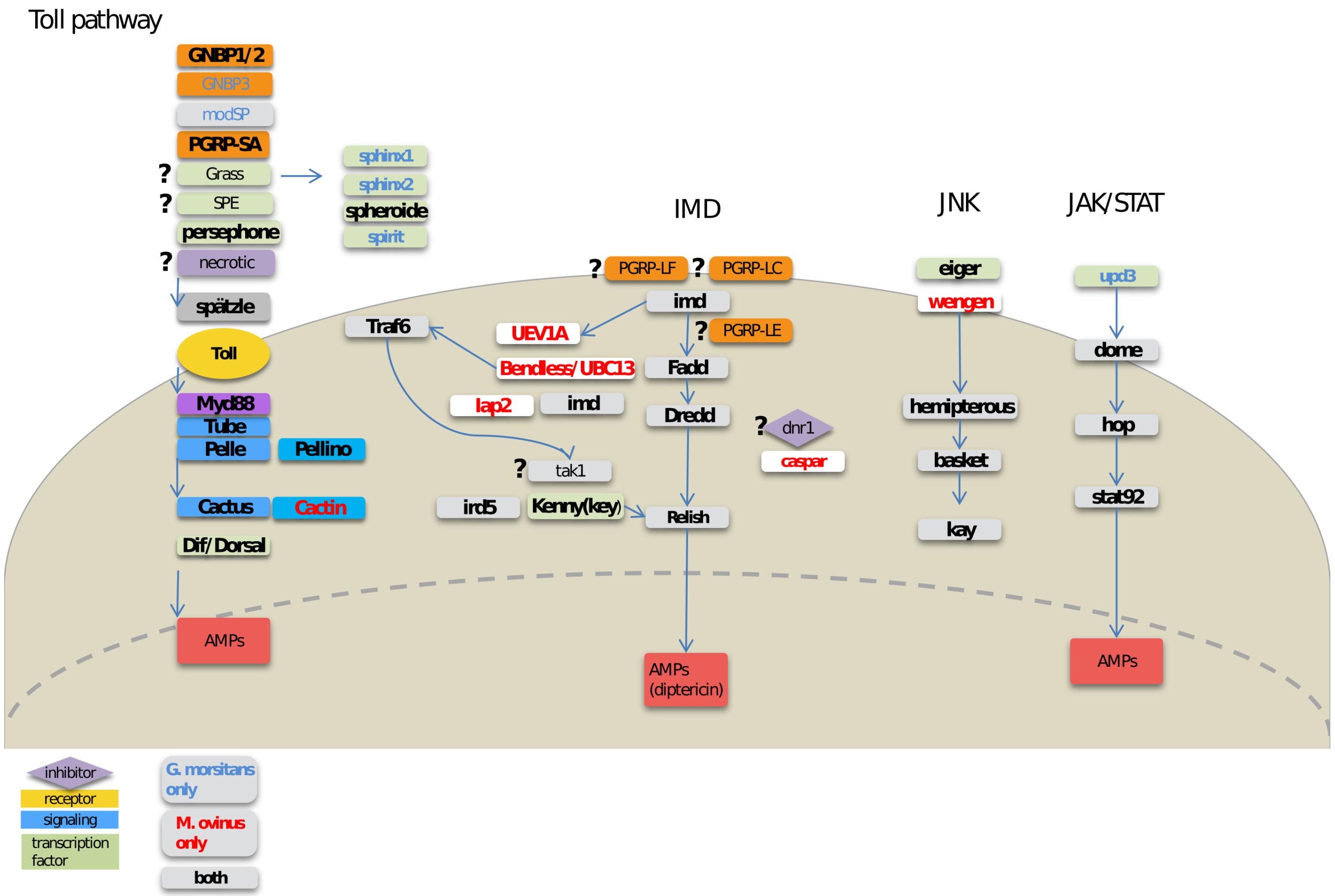


**Supplementary file 3. Immune system genes detected in our reference transcriptome of *Melophagus ovinus*.** Although we show in Fig3a that symbiosis control/maintenance is likely different from tsetse flies, the figure shows that the immune system of *M. ovinus* is almost identical to *G. morsitans*. Genes found to be present/absent in comparison to *Glossina morsitans* are highlighted, but we note that our transcriptome data do not allow for a rigorous analysis of all immune system genes. Gene labels in blue represent genes not detected in the *M. ovinus* transcriptome, but present in *G. morsitans*.

*red: do not correspond to the original query, bold: do correspond, grey: both glossina and drosophila gave the same hit, blue: uncertain.*

| Pathway | Gene name | Gene symbol | Glossina | Melophagus (Blastp) | e value | Drosophila | Melophagus (Blastp) | e value | Melophagus (tblastn) | e value |
|---|---|---|---|---|---|---|---|---|---|---|
| | | | G. morsitans vs. M ovinus blast | | | D. melanogaster vs. M ovinus blast | | | G. morsitans vs. M ovinus blast | |
| Toll pathway | cactin | cactin | - | - | - | cactin | TRINITY_DN33529_c1_g2_i1_p1 | 0 | - | |
| Toll pathway | cactus | cact | GMOY008203 | TRINITY_DN39344_c0_g1_i9_p1 | 5E-65 | cactus | TRINITY_DN39344_c0_g1_i9_p1 | 3.00E-101 | - | |
| Toll pathway | Dorsal-related immunity factor | Dif | GMOY004477 | TRINITY_DN36109_c0_g2_i9_p1 | 0 | DIF | TRINITY_DN36109_c0_g2_i9_p1 | 1.00E-88 | - | |
| Toll pathway | dorsal | dl | GMOY004477 | TRINITY_DN36109_c0_g2_i9_p1 | | dorsal | TRINITY_DN36109_c0_g2_i9_p1 | 0 | - | |
| Toll pathway | Gram-negative bacteria binding protein 3 | GNBP3 | GMOY010453 | TRINITY_DN36993_c2_g1_i1_p1 | 9E-13 | GNBP3 | - | | TRINITY_DN36993_c2_g1_i1_p1 | 0.48 |
| Toll pathway | Gram-negative bacteria binding protein 1/2 | GNBP1/2 | GMOY011181 | - | | | - | | TRINITY_DN38387_c8_g1_i2.p1 | 3.00E-176 |
| Toll pathway | Gram-positive Specific Serine protease | grass | GMOY002008 | TRINITY_DN33120_c3_g1_i3_p1 | 3E-42 | Grass | TRINITY_DN33120_c3_g1_i3_p1 | 2.00E-44 | TRINITY_DN33120_c3_g1_i3_p1 | 5.00E-41 |
| Toll pathway | modular serine protease | modSP | GMOY009437 | TRINITY_DN39489_c2_g3_i5_p1 | 2E-18 | modsp | - | | TRINITY_DN39489_c2_g3_i10 | 1.00E-17 |
| Toll pathway | Myd88 | Myd88 | GMOY005784 | TRINITY_DN32798_c1_g1_i1_p1 | 1E-94 | MyD88 | TRINITY_DN32798_c1_g1_i1_p1 | 2.00E-68 | - | |
| Toll pathway | necrotic | nec | GMOY002444 | TRINITY_DN37921_c2_g1_i3_p1 | 7E-165 | Nec/serpin43Ac | TRINITY_DN37921_c2_g1_i8_p1 | 4.00E-69 | TRINITY_DN37921_c2_g1_i8 | 5.00E-154 |
| Toll pathway | pelle | pll | GMOY000628 | TRINITY_DN33356_c0_g1_i2_p1 | 0 | pelle | TRINITY_DN33356_c0_g1_i2_p1 | 1.00E-137 | - | |
| Toll pathway | Pellino | Pli | GMOY007135 | TRINITY_DN38335_c1_g1_i1_p1 | 0 | pellino | TRINITY_DN38335_c1_g1_i1_p1 | 0 | - | |
| Toll pathway | persephone | psh | GMOY005029 | TRINITY_DN40526_c1_g2_i4_p1 | 1E-102 | persephone | TRINITY_DN40526_c1_g2_i4_p1 | 2.00E-72 | - | |
| Toll pathway | Peptidoglycan recognition protein SA | PGRP-SA | GMOY009549 | TRINITY_DN35844_c2_g1_i1_p1 | 4E-27 | PGRP-SA | TRINITY_DN35844_c2_g1_i1_p1 | 2.00E-70 | - | |
| Toll pathway | Peptidoglycan recognition protein SD | PGRP-SD | GMOY009549 | TRINITY_DN35844_c2_g1_i1_p1 | 4E-27 | PGRP-SD | TRINITY_DN35844_c2_g1_i1_p1 | 6.00E-37 | - | |
| Toll pathway | spatzle | spz | GMOY007932 | TRINITY_DN38025_c3_g4_i2_p1 | 5E-80 | spatzle | TRINITY_DN40507_c1_g1_i4_p1 | 4.00E-28 | - | |
| Toll pathway | Spatzle-Processing Enzyme | SPE | GMOY002535 | TRINITY_DN33120_c3_g1_i3_p1 | 3E-42 | SPE | TRINITY_DN33120_c3_g1_i3_p1 | 7.00E-64 | TRINITY_DN33120_c3_g1_i3 | 3.00E-73 |
| Toll pathway | spheroide | spheroide | GMOY005310 | TRINITY_DN39747_c1_g2_i1_p1 | 4E-61 | spheroide | TRINITY_DN39747_c1_g2_i1_p1 | 2.00E-66 | - | |
| Toll pathway | sphinx1 | sphinx1 | GMOY010124 | TRINITY_DN33925_c1_g1_i1_p1 | 2E-48 | sphinx1/2 | TRINITY_DN32994_c4_g2_i1_p1 | 1.00E-11 | TRINITY_DN33925_c1_g1_i1 | 4.00E-46 |
| Toll pathway | sphinx2 | sphinx2 | GMOY004422 | TRINITY_DN29104_c0_g1_i4_p1 | 3E-25 | sphinx1/2 | - | | TRINITY_DN29104_c0_g1_i4 | 1.00E-24 |
| Toll pathway | Serine Protease Immune Response Integrator | spirit | GMOY008710 | TRINITY_DN33183_c0_g1_i3_p1 | 7E-40 | Spirit | TRINITY_DN29104_c0_g1_i4_p1 | 1.00E-53 | TRINITY_DN33183_c0_g1_i3 | 6.00E-39 |
| Toll pathway | Toll | Tl | GMOY011790 | TRINITY_DN33152_c1_g1_i3_p1 | 0 | Toll-8/Tollo | TRINITY_DN35985_c4_g1_i6_p1 | 1.00E-23 | TRINITY_DN33152_c1_g1_i3 | 0.00E+00 |
| Toll pathway | Toll-7 | Toll-7 | GMOY009848 | TRINITY_DN40715_c2_g1_i7_p1 | 1E-30 | Toll | TRINITY_DN40715_c2_g1_i7_p1 | 3.00E-27 | TRINITY_DN33493_c2_g1_p1 | 0.00E+00 |
| Toll pathway | tube | tub | GMOY007350 | TRINITY_DN38503_c4_g1_i4_p1 | 5E-45 | tube | TRINITY_DN38503_c4_g1_i4_p1 | 6.00E-25 | - | |
| IMD pathway | Caspar | Caspar | - | | | Caspar | TRINITY_DN38960_c4_g2_i1_p1 | 0 | - | |
| IMD pathway | Death related ced-3/Nedd2-like protein | Dredd | GMOY007083 | TRINITY_DN37517_c2_g1_i5_p1 | 7E-141 | dredd | TRINITY_DN37517_c2_g1_i5_p1 | 7.00E-52 | - | |
| IMD pathway | defense repressor 1 | dnr1 | GMOY000299 | TRINITY_DN38691_c2_g1_i4 | 9.00E-08 | | TRINITY_DN38691_c2_g3_i1 | 2.00E-09 | TRINITY_DN34260_c0_g1_i1 | 2.00E-09 |
| IMD pathway | Fas-associated death domain ortholog | Fadd | GMOY007803 | TRINITY_DN35665_c1_g2_i1_p1 | 2E-30 | dFADD/BG4 | TRINITY_DN35665_c1_g2_i1_p1 | 3.00E-31 | - | |
| IMD pathway | Iap2 | | | | | Iap2 | TRINITY_DN35658_c2_g7_i1_p1 | 7.00E-139 | - | |
| IMD pathway | imd | | | | | imd | TRINITY_DN30295_c0_g2_i2_p1 | 1.00E-41 | - | |
| IMD pathway | immune response deficient 5 | ird5 | GMOY007052 | TRINITY_DN39307_c8_g1_i6_p1 | 4E-120 | ird5 | | | - | |
| IMD pathway | kenny | key | GMOY010939 | TRINITY_DN38656_c0_g1_i14_p1 | 3E-97 | kenny | TRINITY_DN38656_c0_g1_i2_p1 | 6.00E-30 | - | |
| IMD pathway | Peptidoglycan recognition protein LC | PGRP-LC | GMOY006094 | TRINITY_DN35844_c2_g1_i1_p1 | 4E-27 | PGRP-LC | | | - | |
| IMD pathway | Peptidoglycan recognition protein LE | PGRP-LE | GMOY006730 | TRINITY_DN34602_c0_g2_i6_p1 | 6E-27 | PRGP-LE | TRINITY_DN35844_c2_g1_i1_p1 | 3.00E-42 | - | |
| IMD pathway | Peptidoglycan recognition protein LF | PGRP-LF | GMOY006730 | TRINITY_DN34602_c0_g2_i6_p1 | 6E-27 | PRGP-LF | TRINITY_DN35844_c2_g1_i1_p1 | 4.00E-39 | - | |
| IMD pathway | relish | | | | | relish | TRINITY_DN38278_c4_g2_i1_p1 | 0 | - | |
| IMD pathway | TGF-beta activated kinase 1 | Tak1 | GMOY005089 | TRINITY_DN28313_c0_g1_i2_p1 | 0 | Tak1 | | | TRINITY_DN38664_c2_g1_i5 | |
| IMD pathway | TNF-receptor-associated factor 6 | Traf6 | GMOY009472 | TRINITY_DN30827_c0_g1_i1_p1 | 0 | Traf6 | | | - | |
| IMD pathway | Bendless/Ubc13 | | | | | Bendless/Ubc13 | TRINITY_DN33596_c0_g1_i1_p1 | 4.00E-108 | - | |
| IMD pathway | UEV1a | | | | | UEV1a | TRINITY_DN36537_c4_g4_i2_p1 | 2.00E-62 | - | |
| JAK/STAT | domeless | dome | GMOY007646 | TRINITY_DN38719_c7_g2_i1_p1 | 0 | Domeless | TRINITY_DN38377_c0_g1_i2_p1 | 9.00E-161 | - | |
| JAK/STAT | hopscotch | hop | GMOY005985 | TRINITY_DN48530_c0_g1_i1_p1 | 8E-49 | hopscotch | TRINITY_DN37993_c2_g2_i2_p1 | 5.00E-45 | - | |
| JAK/STAT | Signal-transducer and activator of transcription protein at 92E | Stat92E | GMOY003394 | TRINITY_DN35508_c7_g1_i6_p1 | 0 | STAT92E | TRINITY_DN35508_c7_g1_i6_p1 | 0 | - | |
| JNK | basket | bsk | GMOY010361 | TRINITY_DN33932_c0_g1_i14_p1 | 0 | basket | TRINITY_DN33932_c0_g1_i14_p1 | 0 | - | |
| JNK | eiger | egr | GMOY005211 | TRINITY_DN38635_c3_g1_i4_p1 | 2E-61 | eiger | TRINITY_DN38635_c3_g1_i4_p1 | 3.00E-50 | - | |
| JNK | hemipterous | hep | GMOY008135 | TRINITY_DN37052_c7_g1_i3_p1 | 1E-119 | hemipterous\| | TRINITY_DN39440_c1_g1_i5_p1 | 1.00E-95 | - | |
| JNK | kayak | kay | GMOY007416 | TRINITY_DN34884_c6_g1_i1_p1 | 8E-95 | kayak | | | - | |
| JNK | wengen | | | | | wengen | TRINITY_DN33427_c6_g1_i1_p1 | 1.00E-60 | - | |
